# Inhibition of extracellular vesicle-encapsulated miRNA produced by estrogen-mediated upregulation of cellular processing suppresses target organ inflammation in a humanized model of systemic lupus erythematosus

**DOI:** 10.1101/2022.11.03.514940

**Authors:** Nicholas A. Young, Emily Schwarz, Rosana A. Mesa, Kyle Jablonski, Lai-Chu Wu, Elisha D.O. Roberson, Wael N. Jarjour

## Abstract

**Background/Purpose:** Distinct, disease-associated intracellular miRNA (miR) expression profiles have been identified from peripheral blood mononuclear cells (PBMCs) of systemic lupus erythematous (SLE) patients. We have previously demonstrated novel estrogenic responses in PBMCs from SLE patients and discovered that estrogen lowers the threshold of immune cell activation to a greater extent in females, including significant upregulation of toll-like receptor (TLR)7 and TLR8 expression. TLR7 and TLR8 bind viral-derived single-stranded RNA to stimulate innate inflammatory responses, but recent studies have shown that miR-21, mir-29a, and miR-29b can also bind and activate these receptors when packaged and secreted in extracellular vesicles (EVs).

**Objective:** The objective of this study was to characterize the estrogen-mediated immunomodulatory effects of distinct EV-encapsulated miR profiles in SLE and evaluate the potential therapeutic approach of miR inhibition in a humanized mouse model.

**Methods:** SLE patients meeting revised ACR guidelines and age/sex-matched healthy controls provided informed consent to participate in this IRB-approved study. Plasma-derived EVs were isolated by differential ultracentrifugation and quantified. PBMCs were isolated from whole blood and cultured in hormone free conditions before stimulation with 17β-estradiol (estrogen; E2). RNA was isolated following E2 stimulation or EV isolation and bulk RNA-sequencing (RNAseq) reads were analyzed. Additionally, PBMCs from active SLE patients were injected into immunodeficient mice to produce chimeras. Prior to transfer, the PBMCs were incubated with liposomal EVs containing complementary locked nucleic acid (LNA) antagonists to miR-21, mir-29a, and miR-29b. After three weeks, blood was collected for both immunophenotyping and cytokine analysis and tissue was harvested for histopathological examination.

**Results:** EVs were found to be increased in the plasma of SLE patients and differentially expressed EV-derived miR profiles were detected compared to healthy controls, including miR-21, mir-29a, and miR-29b. E2 stimulation of PBMCs identified upregulated pathways involved in miR transcription/processing. Specifically, small RNA binding proteins and synthesis enzymes demonstrated significant signaling pathway association and upregulation with E2 treatment. Human immune cell subtypes were successfully recovered from whole blood of chimeric mice at similar levels with and without miR inhibition, but levels of human IL-6, IL-1β, IL-4, and TNF-α were significantly reduced by the LNA antagonists. Moreover, miR antagonists significantly reduced histopathological infiltrates in the small intestine, liver, and kidney, as demonstrated by H&E-stained tissue sections and immunohistochemistry measuring human CD3.

**Conclusion:** These data suggest E2-mediated regulation of miR synthesis and demonstrate distinct EV-derived small RNA signatures representing SLE-associated biomarkers. Targeting upregulated EV-encapsulated miR signaling by antagonizing miRs that may bind to TLR7 and TLR8 reveals a novel therapeutic opportunity to suppress autoimmune-mediated inflammation and pathogenesis in SLE.

## INTRODUCTION

Systemic lupus erythematous (SLE) is a systemic autoimmune disease associated with a myriad of genetic and epigenetic aberrations, environmental triggers, and hormonal influences that affects multiple organs and displays a significant female predominance during reproductive years [1]. Although the adaptive immune response has been investigated extensively in SLE, recent studies suggest that innate immunity may also play a significant role in disease pathogenesis and progression. Lupus nephritis (LN) affects more than 50% of adult and pediatric patients with SLE and is a debilitating clinical manifestation contributing to significant renal-targeted disease morbidity [2]. Despite decades of basic and clinical research investigating SLE, the complex disease pathogenesis remains to be conclusively elucidated and definitively predictive biomarkers of disease activity have yet to be identified. Consequently, patients with SLE have incomplete diagnostic testing available and have limited therapeutic options with targeted agents and biologics.

While extracellular vesicles (EVs) were first reported in the literature half a century ago, their function has only been recently characterized in the last decade. EVs are defined by the International Society of Extracellular Vesicles (ISEV) as membrane-enclosed bodies secreted by cells and can be divided into subpopulations based on size and cellular biogenesis [3]. Exosomes are 50 - 150 nm in diameter and originate from multi-vesicular, intracellular bodies. This study focuses on exosomes, which have the ability to carry highly perishable biocargo in a protected environment once secreted by cells. These biocargo include microRNA (miR) and other small, RNA species. Exosomes facilitate biocargo delivery to a target cell, which provides an important method of cell-to-cell communication. Current evidence is mounting in cancer, infectious disease, and autoimmunity alike for an immunomodulatory role of exosome-derived miRs [3-5]. Moreover, as evidence of diagnostic utility, exosome-associated miR expression patterns can be used to predict drug resistance for patients with multiple myeloma [6].

Small non-coding miRs are about 22 nucleotides in length and are present in animals, plants, and viruses [7-10]. Many miRs have been identified hitherto and each has the potential to target dozens of mRNAs by complimentary binding [11-15]. In this way, the primary, canonical function of miRs has been described as a destabilizer and degrader of mRNA to function as a translational repressor of targets, which ultimately leads to gene silencing [11]. While several recent studies have identified distinct changes in miR expression patterns in SLE [16-19], the identification and validation of novel EV-derived miR signatures may facilitate and guide the next phase of discovery in lupus. Accordingly, studies examining extracellular miR expression and their potential use as a predictor of therapeutic response in lupus patients have been considered. However, a comprehensive profiling of exosome-associated miRs has never been adequately explored.

Recently, we have demonstrated that extracellular miRs are primarily detected in EVs from lupus patients and that EV-encapsulated miR-21 induces both toll-like receptor (TLR)8 and cytokine expression *in vitro* [20]. Furthermore, we conclude that estrogen lowers the threshold of immune cell activation to a greater extent in females and leads to enhanced TLR7 and TLR8 expression [21]. Both TLR7 and TLR8 are X-linked immune system receptors, expressed predominately in macrophages, and stimulate innate inflammatory responses by binding to pathogen-associated, single-stranded (ss)RNA viral sequences. Considering that TLRs serve as an interface between innate and adaptive immunity [22], characterization of this association may contribute to a better understanding of their role in SLE pathogenesis. In addition to the well-characterized pathway of TLR7 and TLR8 activation by binding to viral ssRNAs, recent oncology studies have identified specific miRs capable of activating these receptors when packaged and secreted in EVs [23].

Significantly, this finding characterizes an additional, non-canonical pathway of functionality where EV-delivered miRs can mediate pro-inflammatory signaling in distant cells. While cancer cells have been shown to produce tremendous quantities of exosomes [24], information about EV production and content in the context of autoimmune disease is sparse. Therefore, EV payload and the subsequent downstream effects of miR biocargo on cells still requires extensive investigation in autoimmune diseases, such as SLE, and could translate into clinical diagnostic tests and identify novel pharmacological targets.

This significant, yet unmet need in the SLE diagnostic and therapeutic armamentarium can be addressed by comprehensive examination of bulk RNA species derived from exosomes. Our results identify novel pathways propagated via exosome signaling that represent an opportunity to identify potential therapeutic targets and RNA signatures that can be further evaluated as diagnostic biomarkers. Moreover, our data reveal a potential mechanism of estrogen-mediated miR production and inflammatory signaling in SLE via EVs. These data also suggest that targeting upregulated EV-encapsulated miR signaling by antagonizing miRs that bind to TLR7 and TLR8 may be a novel therapeutic opportunity to suppress autoimmune-mediated inflammation and pathogenesis in SLE. Collectively, these novel inflammatory mechanisms propagated via EV signaling present an opportunity to gain invaluable insight into SLE pathogenesis. The elucidation of EV-based pathways by miR profiling can unveil targets that could be translated into long-overdue therapeutics or more sensitive diagnostic tests.

## METHODS

### Human Samples and collection

Lupus patients were recruited from The Ohio State University Wexner Medical Center (OSUWMC) Lupus Clinic and healthy controls from the local community. Lupus patients met the revised criteria of the American College of Rheumatology for SLE [25]. All participants in the study provided informed consent and the research was conducted under an IRB approved protocol at OSUWMC. Blood samples were collected into EDTA containing tubes. Plasma was isolated by centrifugation, aliquoted, and frozen at -80^°^C. PBMCs were isolated by density gradient centrifugation over Ficoll as previously described [26].

### Hormone treatment

After PBMCs isolation, cells were cultured in hormone free conditions as previously described [26] using RPMI 1640 (Life Technologies) and 5% charcoal stripped fetal bovine serum (Life Technologies). After resting overnight, cells were treated with 10 nM of 17β-estradiol (E2; Sigma-Aldrich) and collected after a 36 hr incubation according to previously established protocols [21].

### RNA isolation and RT-qPCR

Cellular RNA was isolated, quantitated, synthesized to cDNA, and used for quantitative (q)PCR following previously described methods [14]. Briefly, RNA was isolated from PBMCs using the RNeasy Mini Kit (Qiagen Sciences) and from whole blood samples using the Paxgene Blood RNA Kit (PreAnalytix; Qiagen) according to manufacturer’s protocol. For analysis of RNA in EVs, total RNA was isolated according to manufacturer’s protocol using the MagMAX mirVana total RNA isolation kit (Thermo Fisher Scientific). All RNA isolations were quantitated using a NanoDrop 1000 spectrophotometer (NanoDrop Products) and were processed in the presence of chemical RNase inhibitor, RNAsecure (Thermo Fisher Scientific). cDNA was synthesized using the High Capacity cDNA Reverse Transcription Kit (Applied Biosystems) following manufacturer’s protocol and qPCR was performed using the TaqMan system (Applied Biosystems) with cDNA and gene-specific primers according to manufacturer’s protocol. All samples were run on the ABI Prism 7900HT Sequence Detection System (Applied Biosystems) and normalized to 18sRNA as an internal positive control. Results were analyzed using the 2^-ΔΔCt^ method [27].

### Exosome isolation and quantitation

Exosome isolation was carried out using the Cold Spring Harbor ultracentrifugation protocol [28]. Briefly, biospecimens (blood or urine) were centrifuged to remove any present cells and supernatants were then centrifuged to remove any remaining cell debris. The resulting supernatants were collected, transferred to new tubes, and centrifuged at 100,000 x g for 70 min at 4°C. The pelleted exosomes were resuspended in PBS, transferred to new centrifuge tubes, and centrifuged again at 100,000 x g for 70 min at 4°C to remove any contaminating protein aggregates that may have sedimented with the exosome pellet. The subsequent pellet containing purified exosomes was stored at -80°C prior to further processing.

The NanoSight LM10 nanoparticle tracking analysis (NTA) system was used to measure the rate of Brownian motion of particles to determine the size and concentration of EVs according to manufacturer’s instructions (NanoSight). Briefly, samples were injected into the cubicle and the size of particles was obtained by determining the motion in fluid passing across the chamber. The particles/mL were measured and acquired from both experimental and control samples for comparison using area under the curve calculations obtained via the linear trapezoidal analysis method. Plasma was added to enzyme-linked immunosorbent assay (ELISA) ExoTEST plates (Hansa BioMed) according to manufacturer’s protocol and exosome concentrations were measured.

### RNA library preparation and analysis of high-throughput sequencing data (exosomes)

Following exosome isolation, RNA concentrations were determined using the Qubit high-sensitivity RNA assay and small RNA libraries were created using the Clontech SMARTer smRNA-Seq preparation kit according to manufacturer instructions (Takara). Small RNA inserts were further enriched using AMPure bead selection (Beckman Coulter) prior to quantifying with the Qubit DNA high-sensitivity kit. Equimolar amounts of each sample library were pooled to make a single collection for each sequencing run to limit batch effects. Libraries were sequenced on an Illumina HiSeq3000 with 50 base-pair, single-end reads. Raw sequencing read files were modified with cutadapt to remove sequencing adapters and non-template bases added during library preparation. miRs were counted by aligning (RNA-STAR) either to the whole GRCh38 human genome or the annotated human miRs (miRBase).

### RNA library preparation and analysis of high-throughput sequencing data (PBMCs)

RNA was isolated from PBMCs according to manufacturer’s protocol using the RNeasy Mini Kit (Qiagen Sciences) after stimulation with estrogen in culture. Cell pellets were collected at 36 hrs and RNA sequencing for mRNA and miRNA transcriptional targets was performed through the Nucleic Acid Shared Resource (NASR) at OSUWMC. Targets with low counts were removed via edgeR::filterByExpr; this uses embedded filtering for sequencing depth and experimental design considerations. After filtering, 24,511 genomic targets were included in the mRNA analysis and 1,599 were included in the miRNA analysis, which correlates to 85% and 92% of the total, respectively. Normalization was done through the trimmed mean of M-values method and indicated that the filtering was adequate for downstream comparative evaluation. Using these data, results were normalized to untreated values for each subject individually to account for subject-to-subject variability in the datasets. Ingenuity® pathway analysis (IPA) software was then used to filter the genes associated with estrogen treatment compared to untreated controls and selected for those known to be associated with immunological disorders or immune function, as defined by Ingenuity IPA Systems and according to our previously established analysis methods [21].

### Mice and sample size determination

4-week old NOD.Cg*-Prkdcscid Il2rgtm1Wjl*/SzJ (NSG) and NOD.Cg-*Rag1*^*tm1Mom*^ *Il2rg*^*tm1Wjl*^/SzJ (NRG) mice were obtained from The Jackson Laboratories. Mouse maintenance and protocol 2017A00000032 was approved by the Institutional Animal Care and Use Committee at OSUWMC with adherence and recognition of ARRIVE guidelines. The animal facility was maintained at 22-23°C and between 30-50% relative humidity with a 12-hour light/dark cycle. Chow and water were available ad libitum. The sample sizes were based on calculations using reference data from previous studies with type one error of 0.05 and power of 0.8. To verify reproducibility and rigor of the data, experiments were repeated in accordance with NIH reproducibility guidelines. The number of mice used in each experiment was designed to provide a statistically significant result with a minimum number of animals. Because of possible sexual dimorphism for immune response, male and female mice were studied separately.

### Adoptive Transfers

Freshly isolated human PBMC preparations were washed in PBS and counted using a hemocytometer with trypan blue to ensure cell viability. All samples were kept separate and not pooled before injections. Prior to adoptive transfer into mice, PBMCs were treated with 100 nM total of human miRCURY (Exiqon) locked nucleic acid (LNA) inhibitors or equimolar negative control (scrambled LNA): miR antagonists included miR21 CAACATCAGTCTGATAAGCT; miR29a AACCGATTTCAGATGGTGCT; and miR29b ACTGATTTCAAATGGTGCT. LNA-anti-miRs were combined with DOTAP liposomal transfection reagent according to manufacturer’s protocol and incubated approximately 10 hrs. PBMCs were injected intraperitoneally into 8-week old NSG or NRG mice (5.0 × 10^6^ cells/mouse); at least 2-3 mice were injected from each individual human sample. Mice were monitored every other day, including weights and physical signs of disease progression, and sacrificed three weeks after adoptive transfer for blood and tissue collection as described below.

### Cytokine measurement

Murine serum was prepared from whole blood collected via subclavian artery at the experimental endpoint. Serum samples were analyzed by ELISA using the V-PLEX Pro-inflammatory Panel 1 mouse kit (Meso Scale Diagnostics) according to the manufacturer’s protocol and data was analyzed using Microsoft Excel (v2016), as previously described [29]. All reported results were above the limit of detection (LOD) of the assay.

### Tissue collection and histopathology

Tissues were resected from each mouse and flash frozen in liquid nitrogen followed by cryosectioning or immediately fixed by immersion in neutral buffered 10% formalin and then processed into paraffin (FFPE). Serial histological sections were stained with H&E (Leica Microsystems) following the manufacturer’s protocol, or labeled by immunohistochemistry (IHC) as detailed previously [29]. Histological slides were digitally scanned for downstream quantitative image analysis.

H&E-stained FFPE tissue sections were subjected to blinded histopathological analysis using the 10x objective of a bright-field light microscope with predefined criteria [29, 30]. Scanned slide images were analyzed by Aperio ImageScope digital analysis software (v9.1) as detailed previously to determine positive staining and lymphocyte localization by IHC [30, 31]. Specifically, Aperio’s positive pixel count algorithm was run to quantify the extent of positive staining and lymphocyte localization using calibrated hue, saturation, and intensity values following previously described methods of computer-assisted image analysis [32]. To define a mean positive pixel intensity value for each IHC slide, 10 measurements of identical total surface area of tissue were quantitated. All digital analysis was confirmed by manual slide interpretations using the 10x objective of a bright-field light microscope.

### Flow Cytometry

Blood was collected from chimeric mice by submandibular bleeding and leukocytes were purified for flow cytometry using ammonium chloride solution for red blood cell lysis. Cells were subsequently plated and blocked with anti-mouse FcR antibody (αCD16/CD32, BioLegend) in FACS buffer (PBS with 2% BSA and 1mM EDTA). Cells were then surface-stained with antibodies for anti-human CD3 (eBioscience), CD4 (Immunotech), CD8 (Caltag Laboratories), CD20 (eBioscience), CD14 (eBioscience), or CD56 (eBioscience) following manufacturer’s protocol. Data were collected on the BD LSRII Flow Cytometer (BD Biosciences) platform and exported for analysis via FlowJo (v.9.0; Treestar, Inc).

### Statistics

Statistical analysis was performed by one-way ANOVA followed by Tukey’s post hoc test for multiple comparisons (Graph Pad Prism software v8.3.0) or by paired, two-tailed, Student *t*-tests (Microsoft Excel v2016). All numerical datasets were expressed as mean values with standard error of the mean (SEM) indicated using Graph Pad Prism or as mean values ± standard deviation using Microsoft Excel. Data was considered statistically significant if p ≤ 0.05.

## RESULTS

### RNA expression profiles from extracellular vesicles are specific to biological sources and differentially regulated in active lupus patients

To evaluate whether enhanced exosome production and/or differential miR content within exosomes can contribute to the heightened inflammatory state observed in SLE, we isolated exosomes from the plasma or urine of healthy controls or active LN patients. Exosome isolation was carried out using the differential ultracentrifugation protocol, as outlined in the Cold Spring Harbor protocol [28, 33]. This method remains a gold-standard in exosome studies and has been approved as a valid methodology by the ISEV [3]. Following exosome isolation, the NTA system was used to measure EV size and concentration. While initial plasma input contained EVs of various size, the ultracentrifugation isolation protocol yielded purified EVs consistent with exosome size, 100-200 nm diameter (**Figure 1A**). Subsequently, small RNAs were quantitated via Agilent bioanalyzer and normalized to a starting concentration prior to RNAseq. The content of these exosomes demonstrated that they are indeed enriched for RNA species corresponding to typical miR size, approximately 22 nucleotides in length (**Figure 1B**). Each sample was eluted and small RNAseq of at least 1 ng total RNA input was performed. Using the current version of miRbase, even at shallow sequencing depth and without further insert size optimization, we identified at least 94 known human miRs with at least 5 reads in all samples (data not shown).

**Figure 1.**
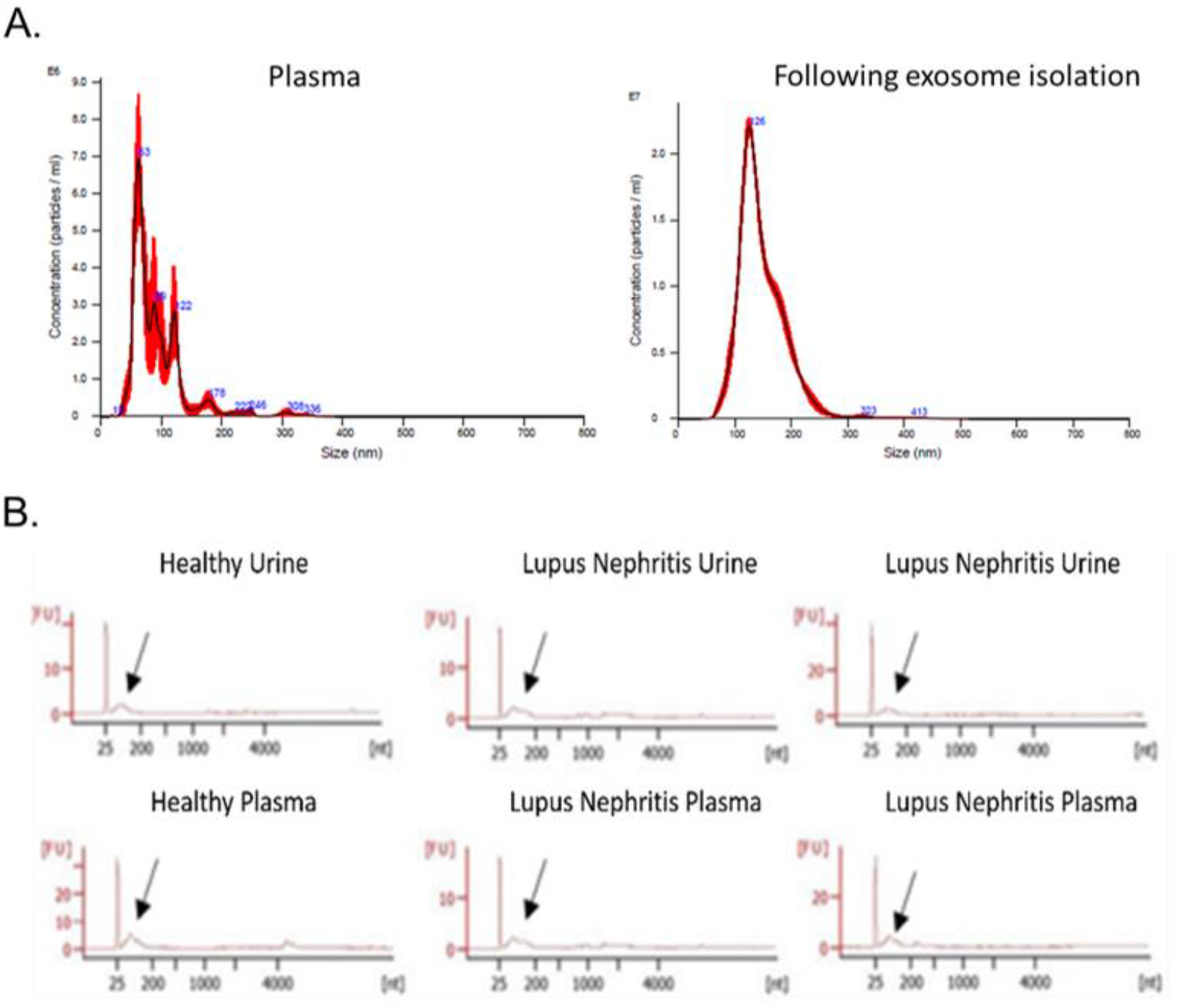
Extracellular vesicle characterization demonstrates representative exosome size and enriched miR fractions. **(A)** Following EV isolation by ultracentrifugation, biological samples were analyzed by nanoparticle tracking. **(B)** Bioanalyzer traces showing that the miR fraction is detectable from exosome isolations. Representative bioanalyzer traces showing miR fraction detectable from both urine and plasma samples derived from LN patients and healthy subject exosome samples. Arrows indicate small RNA (lncRNA and miR).

While exact starting volumes of plasma varied from sample to sample, RNA yields were validated from exosomes isolated from both plasma and urine despite differences in initial volumes; thus providing sufficient samples for multiple library attempts from every isolation. Principal components analysis (PCA) of the annotated small RNA species distinctly separated urine-derived exosomes from plasma-derived exosomes in healthy control samples (**Figure 2A**). These results demonstrate that biological source can differentiate exosome-derived miR profiles. Similarly, PCA of bulk small RNA profiles from plasma exosomes also demonstrated a strong correlation between groups when comparing active LN patients to healthy controls (**Figure 2B**). Furthermore, normalization of expression levels and differential expression corrections yielded significant differences in RNA targets derived from plasma exosomes of active LN patients relative to controls. The greatest differential expression results included vault (v)RNAs (*VTRNA1-1* and *VTRNA1-2*). All vRNA family members share the same chromosomal location (5q31.3) and were each downregulated >20-fold (q-value 8.51E-08; q-value 9.89E-07). The extent and significant differences in bulk RNA signatures are markedly observed when displayed in a volcano plot (**Figure 2C**). The differentiation of body fluid samples by PCA and unique plasma-derived RNA expression profiles indicate that lupus patient exosomes are distinct from healthy controls.

**Figure 2.**
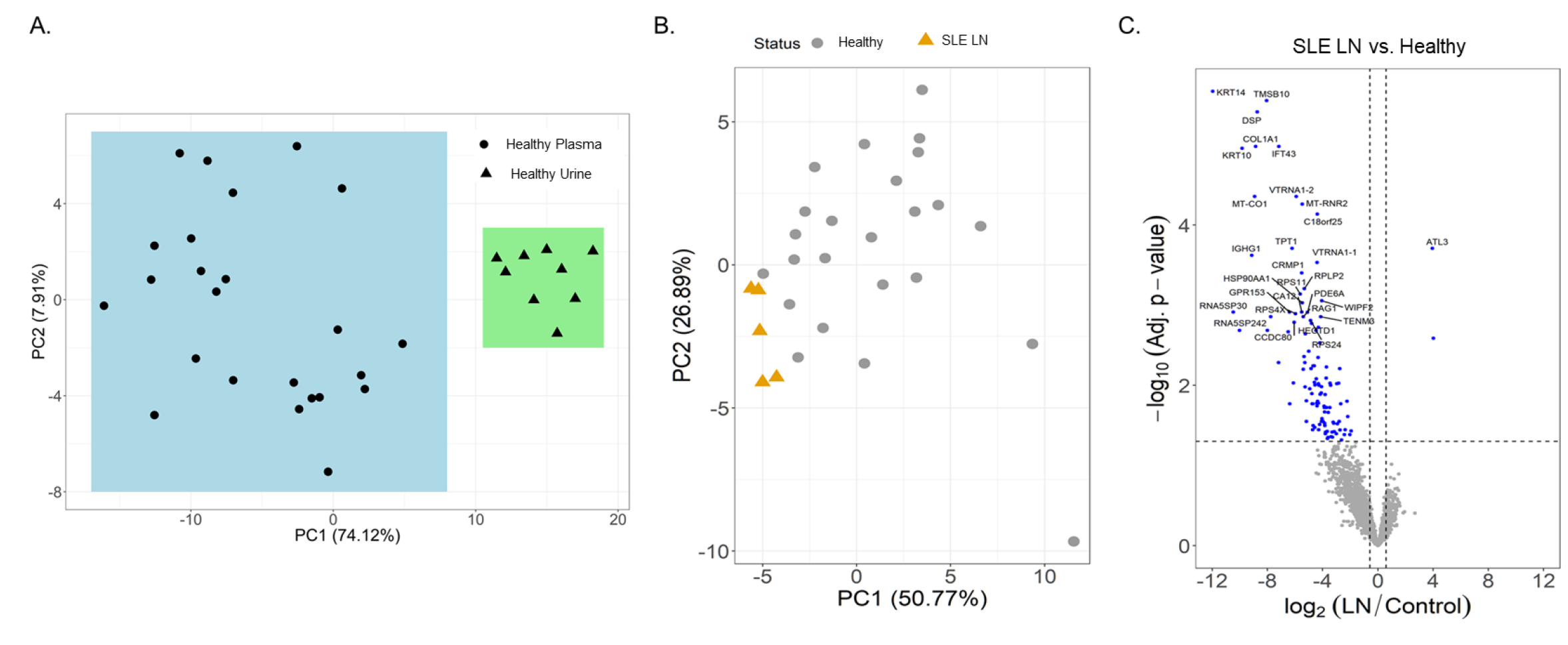
RNA expression profiles from plasma-derived extracellular vesicles (EVs) are distinct from urine and differentially regulated in patients with lupus nephritis (LN). **(A)** EVs were isolated by ultracentrifugation from healthy human urine (N = 10) and plasma (N = 25) sources. **(B)** Plasma-derived EVs were isolated by ultracentrifugation from LN patients (N= 5) and healthy controls (N = 25). **(A-B)** RNA was isolated from EVs for comprehensive, bulk RNA-sequencing analysis for detectable small RNA sequences. Reads were aligned and principal component analysis was performed to evaluate grouping of the datasets. **(C)** A volcano plot was made from the data generated in **(B)**.

### Extracellular vesicles and EV-encapsulated miRs that bind and activate TLR7 and TLR8 are upregulated in SLE patients with lupus nephritis

To further explore the association of exosome-encapsulated RNA biocargo and disease activity, we isolated plasma from SLE patients (including inactive/active and with/without renal involvement) and compared exosome concentrations to age/sex-matched healthy controls. SLE exosome levels were higher than healthy controls both with and without active disease or renal involvement (**Figure 3A**). Specifically, exosome concentrations of active SLE patients were 2-fold greater when compared to healthy controls and statistically significant. Since exosome concentrations are higher in SLE on average, normalized RNA values will not physiologically reflect relative levels when comparing SLE to healthy control samples. Consequently, while the exosome-derived small RNA profiling above was valuable to establish differential content, individual target expression levels from total RNA are needed to evaluate potential functional impact *in vivo*. To this accord, specific exosome-derived miR targets were selected based on published data and included miR-21 [23], miR-29a [23], and miR-29b [34]; each has been shown to be packaged into exosomes and signal through recipient cells via TLR7 or TLR8. While no difference was observed in Let7a, expression of miR-21, miR-29a, and miR-29b was significantly elevated in active SLE patients with LN relative to age/sex-matched healthy controls (**Figure 3B**). Considering that these exosome-encapsulated miRs can activate TLR7 or TLR8 and that immunomodulatory exosomes are upregulated in SLE, combinational antagonism of miR-21, miR-29a, and miR-29b represents a novel therapeutic strategy to evaluate.

**Figure 3.**
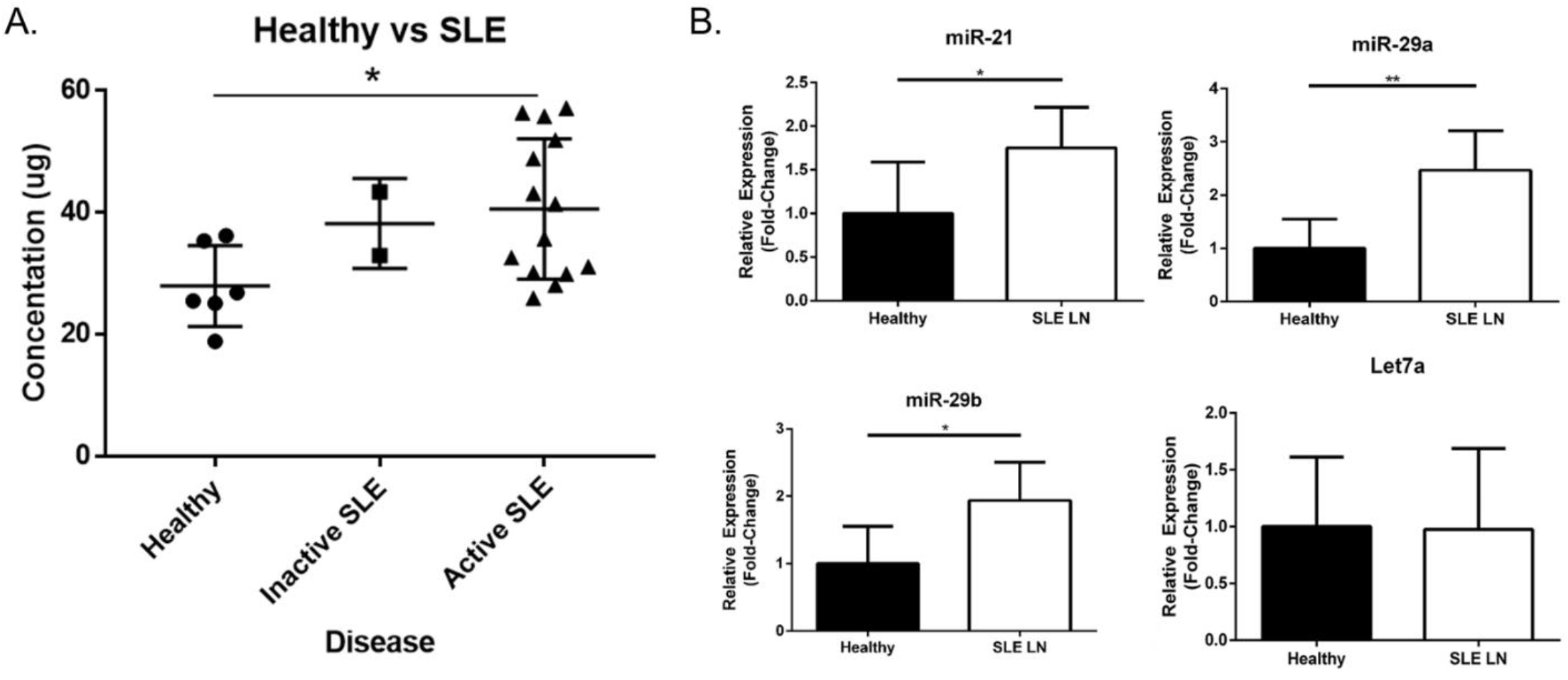
Extracellular vesicles and EV-encapsulated miRs that bind and activate TLR7 and TLR8 are upregulated in SLE patients. **(A)** Plasma derived EVs were isolated from healthy volunteers (N = 6) and SLE patients (N = 16; 14 with active disease and 2 inactive) by ultracentrifugation and quantified by an ELISA assay. **(B)** RNA was extracted from EVs isolated from healthy volunteers (N = 6) and active SLE patients (N = 6). Expression of miR-21, miR29a, miR-29b, and Let7a was measured by RT-PCR analysis. Data was normalized to RNU-44 internal control expression and shown as a relative fold-change. Values are the mean ± SEM with indicated p values calculated via paired, two-tailed, Student’s t tests. *p ≤ 0.05; **p ≤ 0.01.

### RNA sequencing of PBMCs reveals estrogen-mediated upregulation of miR processing machinery

Studies examining intracellular miR expression in PBMCs of patients with SLE have identified distinct disease-associated changes [35]. Furthermore, our previous work has demonstrated that estrogen can significantly upregulate both TLR7 and TLR8 expression and lower the threshold of immune cell activation more extensively in females [21]. To examine the effects of estrogen on miR processing and production, human PBMCs isolated from whole blood were cultured in hormone free conditions overnight before stimulation with 10 nM of 17β-estradiol (estrogen; E2). Total RNA was isolated from cell lysates and RNA libraries were prepared for RNAseq; E2-treated samples were compared and normalized to untreated controls for each individual donor. Comprehensive signaling and network pathway analysis of RNAseq data predicted upregulation of estrogen-dependent breast cancer signaling by canonical pathway overlay and estrogen receptor upregulation by overlap upstream analysis (data not shown). Additionally, further overlay analysis identified upregulated pathways that included several involved in miR transcription/processing (**Figure 4A**). Specifically, argonaute-2 (AGO2) showed an IPA overlap p-value of 0.006, which resulted from upregulation of AGO2 mRNA expression by 1.3-fold with E2 treatment and significant downregulation of FOS, miR-127, miR-34, and miR-27. Moreover, Dicer1 mRNA (a Ribonuclease III enzyme) was stimulated 1.3-fold with estrogen treatment and an IPA overlap p-value of 0.001 was observed that resulted from miR-34, miR-196, and miR-218 suppression. Estrogen-induced mRNA expression in expected genes based on our previous studies as well as other genes involved in intracellular miR processing were also observed (**Figure 4B**). Specifically, Drosha RNase III mRNA was induced 1.2-fold with estrogen treatment and several members of the RNA polymerase III family (POLR3A, POLR3D, and POLR3D) were upregulated 1.3-fold, 1.4-fold, and 1.3-fold, respectively. In concordance with our previous studies [21, 26, 36], estrogen-mediated upregulation of STAT1 (1.4-fold), ESR1 (1.3-fold), ESR2 (1.2-fold), TLR8 (1.5-fold), STAT4 (1.3-fold), and TLR7 (1.3-fold) were also demonstrated.

**Figure 4.**
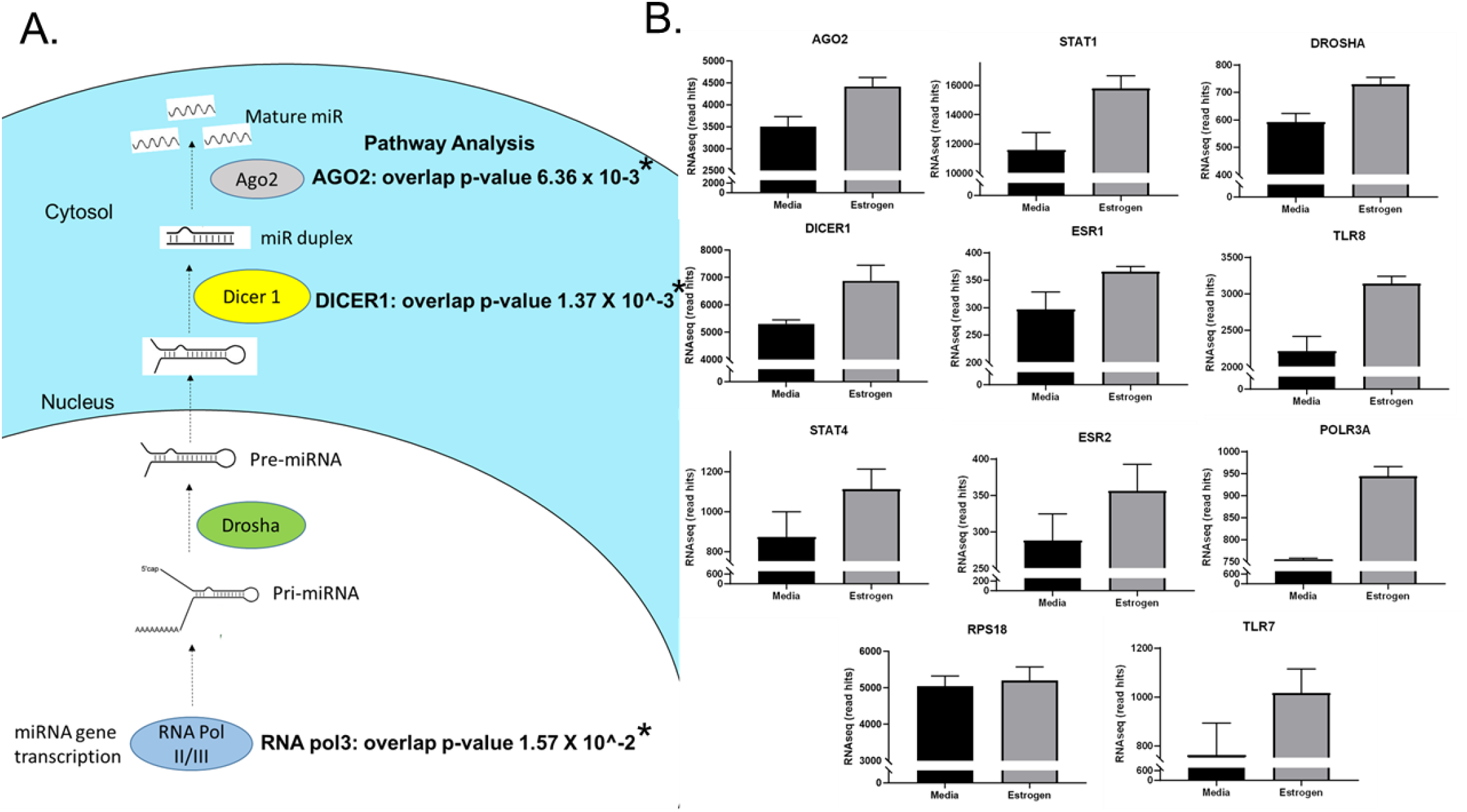
Estrogen induces intracellular miR processing pathways in human PBMCs. Human PBMCs were isolated from healthy volunteers (N = 6) and stimulated with 17β-estradiol (estrogen; E2). RNA was isolated for RNA-sequencing analysis for detectable miR and mRNA sequences. Reads were aligned and analyzed by Ingenuity Pathway Analysis tools examining comprehensive effects of mRNA and miR expression on cellular signaling pathway activation. **(A)** Overlapping p-values from Ingenuity analysis are indicated for select proteins involved in miR synthesis. **(B)** RNA-sequencing read hits are indicated for miR synthesis pathway mRNAs and other known estrogen-regulated genes. *All values of p ≤ 0.05 considered statistically significant.

### Inhibition of disease progression in a human-mouse chimeric model of inflammation by antagonizing miRs that bind TLR7 and TLR8

Our previous data has shown that liposome-encapsulated miR-21 can indeed signal through TLR8 to induce TLR8 expression, pro-inflammatory cytokine signaling, and additional EV secretion in human macrophages and primary PBMC cultures [36]. Furthermore, we have also demonstrated that locked nucleic acid (LNA) modification can be used as an antagonist to block the induction of TLR8 expression by endogenous EVs [20]. To block exosome-encapsulated, miR-induced inflammation via miR-21, miR-29a, and miR-29b, human PBMCs derived from active LN patients were treated with a cocktail of LNA miR antagonists (LNA cocktail) or equimolar of LNA control (miRScramble) prior to adoptive transfer into immunodeficient murine recipients. LNAs contain a modified ribose moiety and were used here to antagonize miR functionality *in vivo*. They are significantly more resistant to enzymatic degradation than RNA and have been shown to persist *in vivo* for over a week [37]. Adoptively transferred PBMCs were engrafted into immunodeficient mouse recipients for three weeks; our previous work recapitulating autoimmune disease in the NSG mouse model demonstrated that this period of time was optimal for histopathological observation of target organ inflammation without confounding graft versus host reaction [31]. Human PBMC engraftment to establish human-mouse chimeras in immunodeficient murine backgrounds all have associated limitations and considerations [38]. Consequently, both NRG and NSG immunodeficient mice were used for human-mouse chimeric modeling and demonstrated similar results in this study. Successful PBMC engraftment was observed in both models and with both LNA treatment conditions, as demonstrated by similar levels human T-cells (CD4+ and CD8+), B-cells, monocytes, and NK cells recovered from whole blood of all chimeric mice (**Figure 5**). Although similar levels of PBMC engraftment were observed, disease progression was indicated in mice treated with LNA control; while weights of uninjected immunodeficient mice were similar trending longitudinally to LNA cocktail, LNA control-treated mice showed a notable loss of body mass at the three week timepoint (**Figure 6**).

**Figure 5.**
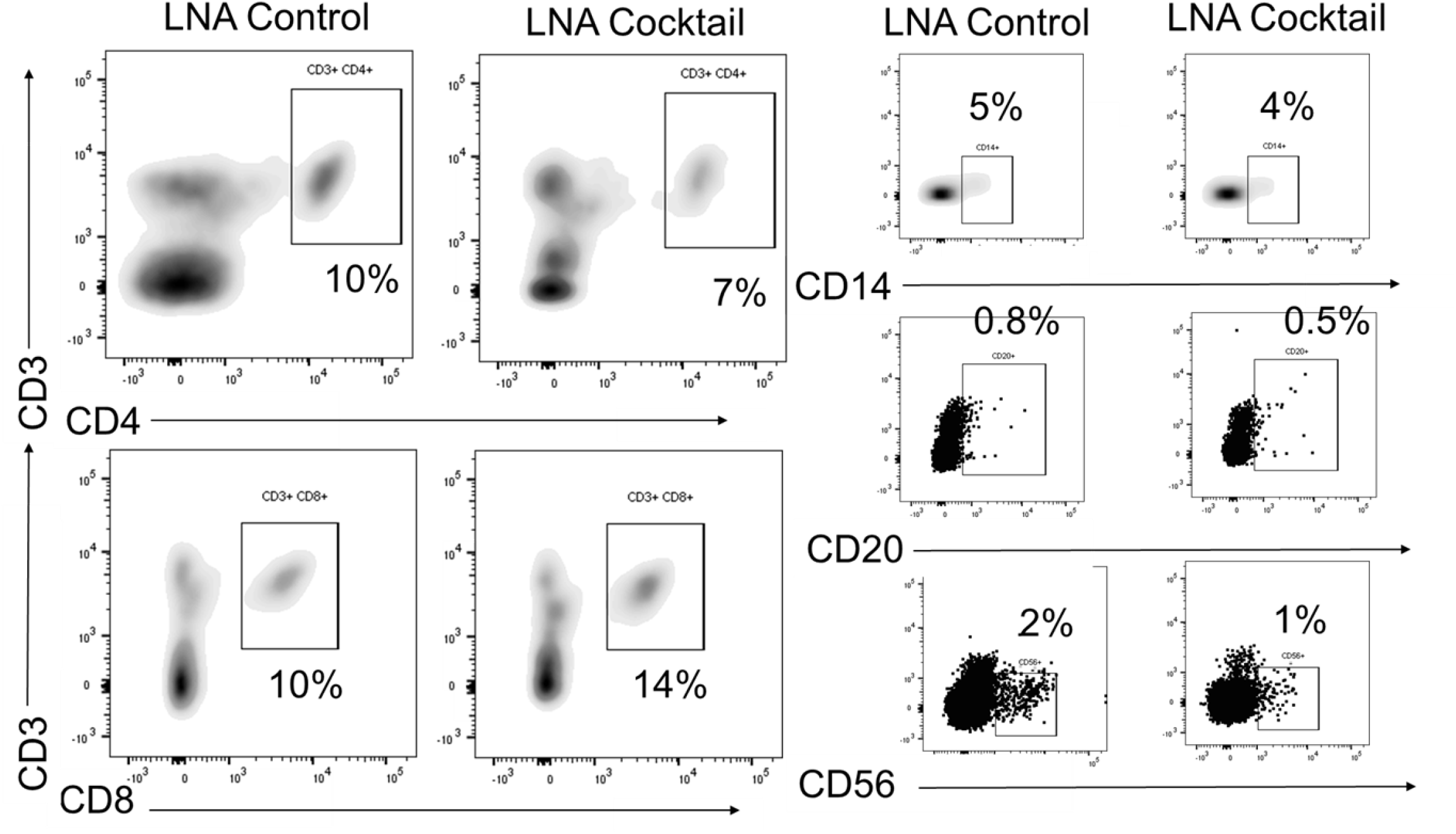
Reconstitution of human PBMCs is observed with control or miR inhibition. Whole blood was analyzed by flow cytometry in chimeric mice following treatment with LNA cocktail or LNA control. Immune cell subtype distribution was measured using the indicated markers, as detailed in the methods section.

**Figure 6.**
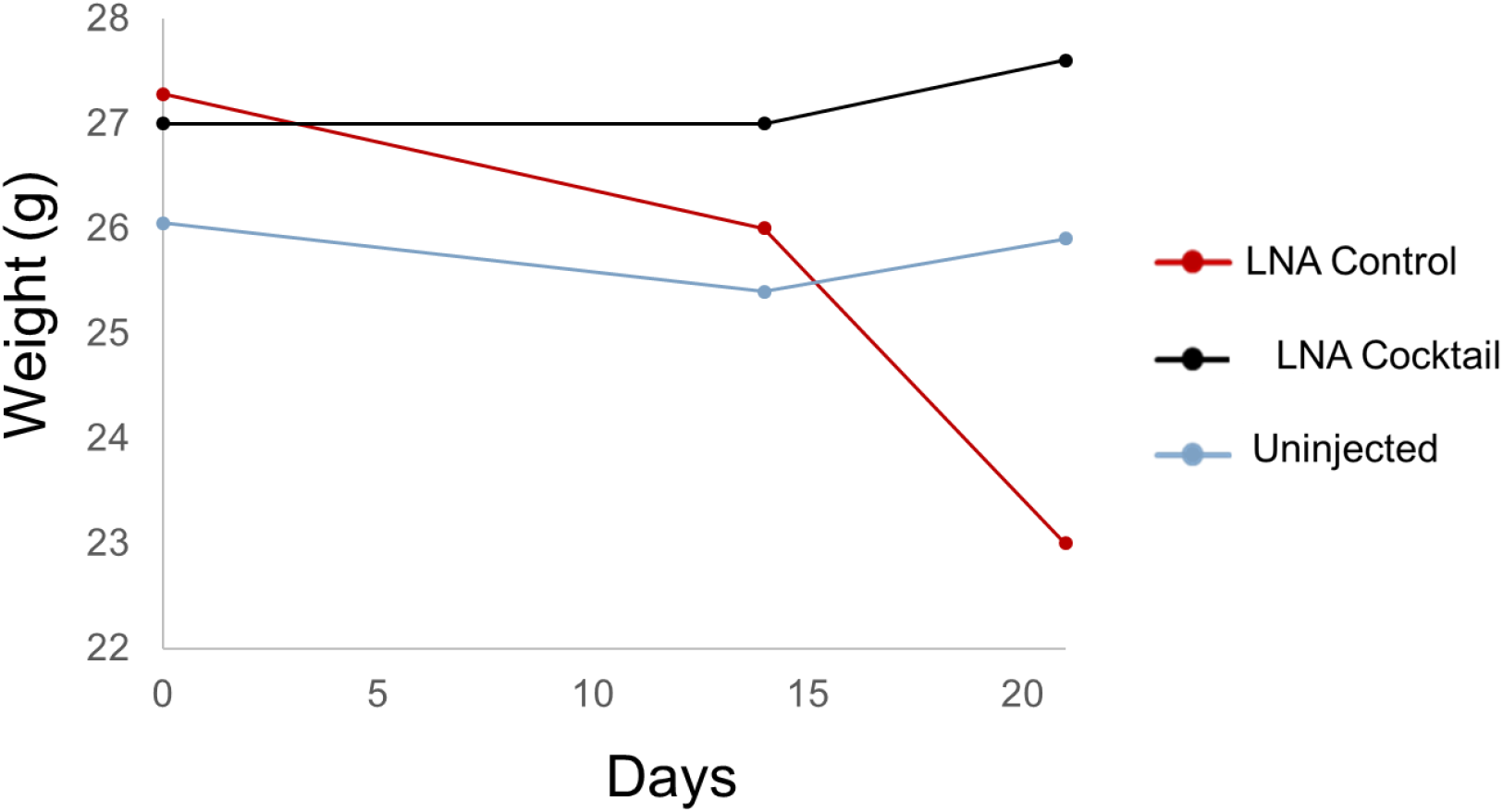
Expected weight-loss prevented with inhibition of miR targets in humanized murine model. Weights were recorded over time in chimeric mice treated with locked nucleic acid miR antagonists (LNA cocktail) targeting miR-21, miR-29a, and miR-29b for comparison to a nonsense control treatment (LNA control).

To assess the effects of antagonizing miR-21, miR-29a, and miR-29b with LNA inhibitors for comparison to LNA control, serum and tissue samples were collected from NSG and NRG mice. Serum was analyzed for proinflammatory cytokine expression to measure the systemic response at three weeks post-adoptive transfer. Treatment with the LNA inhibitor cocktail targeting miR-21, miR-29a, and miR-29b resulted in significant decreases in IFN-γ by 40% (p = 0.0068), TNF-α by 67% (p = 6.73 × 10^−5^), IL-6 by 70% (p = 2.25 × 10^−7^), IL-2 by 47% (p = 0.0085), and IL-4 by 63% (p = 0.0043) relative to PBMCs treated with LNA controls (**Figure 7**). While little to no inflammation was observed in the skin and ear, histopathological examination by H&E showed a robust inflammatory response in the small intestine, liver, and kidney with control treatment, which was markedly reduced with miR inhibition via LNA cocktail treatment (**Figure 8**).

**Figure 7.**
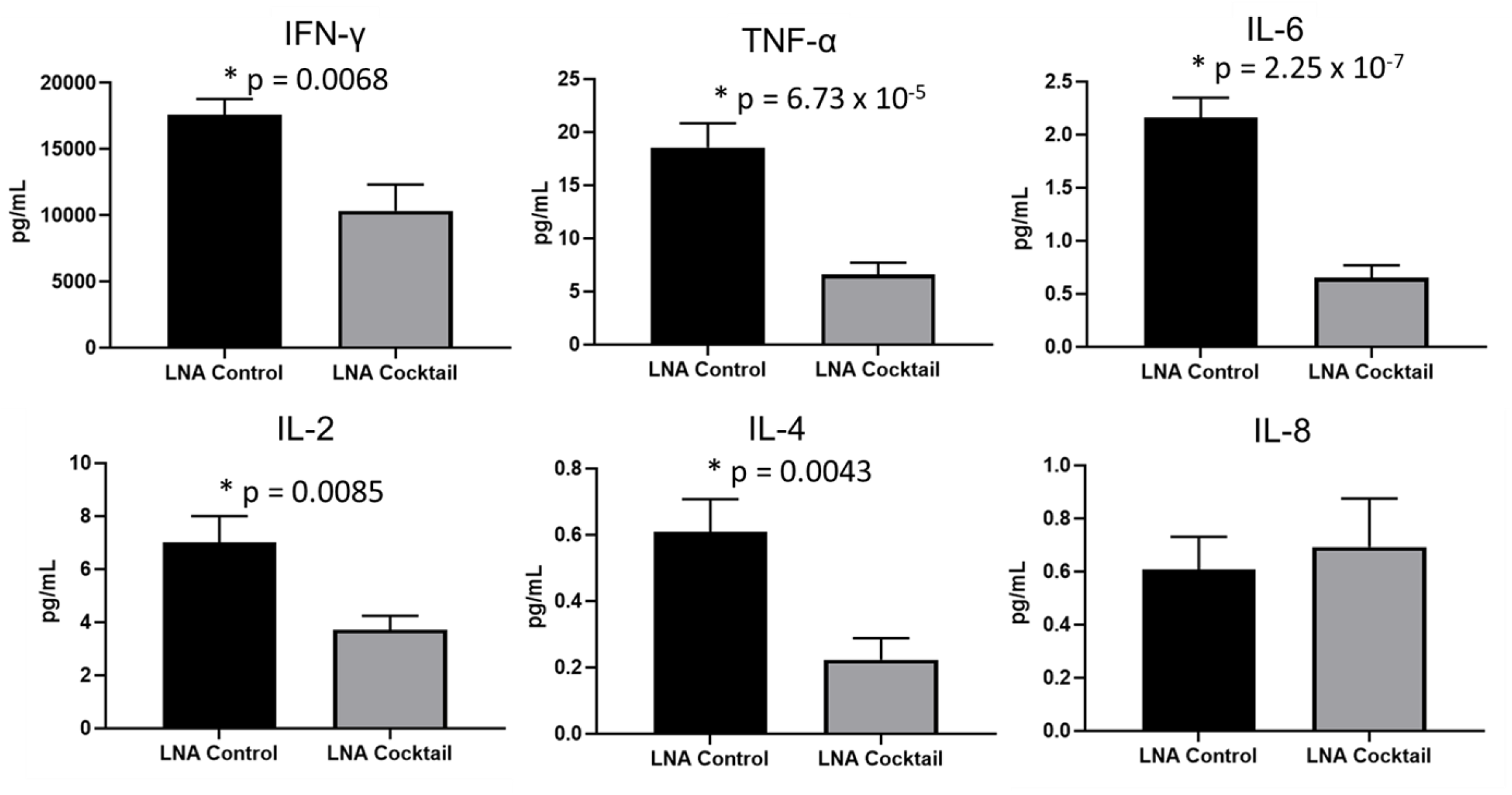
Proinflammatory cytokine expression is reduced in chimeric mice treated with miR antagonists. PBMCs from active SLE patients (N = 4) were transfected *ex vivo* with a cocktail of locked nucleic acid miR antagonists (LNA cocktail) targeting miR-21, miR-29a, and miR-29b, or a nonsense control (LNA control). Serum proinflammatory cytokine expression was measured in chimeric mice (N = 10) after 21 days by ELISA. Values are the mean ± SEM with indicated p values calculated via paired, two-tailed, Student’s t tests. *p ≤ 0.05; **p ≤ 0.01.

**Figure 8.**
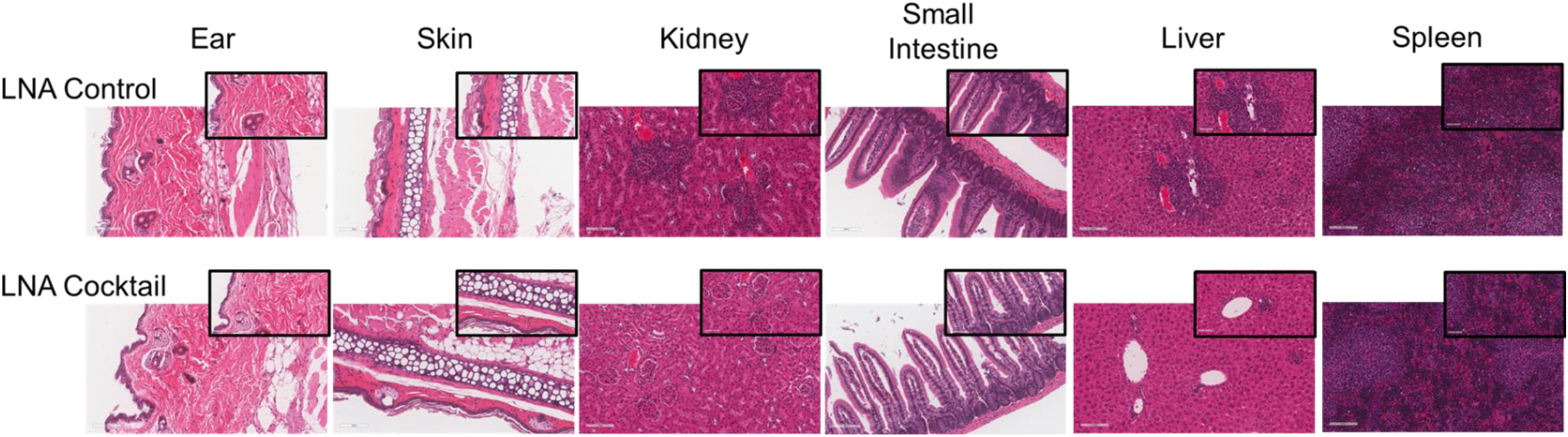
Inhibition of target organ inflammation by antagonizing miRs that bind TLR7 and TLR8 suppresses inflammation in a novel human-mouse chimeric model. PBMCs from active SLE patients were adoptively transferred into immunodeficient mice to produce chimeras. For each SLE patient (N = 4), 2 or 3 immunodeficient mice were used, depending on cell yield (N = 10 total). To block miR-induced inflammation, PBMCs were transfected *ex vivo* with a cocktail of locked nucleic acid miR antagonists (LNA cocktail) targeting miR-21, miR-29a, and miR-29b, or a nonsense control (LNA control) prior to injection. 21 days after adoptive transfer, tissues were collected and processed for H&E staining. Large panel images taken at 20X magnification (scale bar representing 100 μm) and insets taken at 40X (scale bar representing 50 μm).

Specifically, renal inflammation was characterized by infiltrates into the glomerulus leading to abnormal histological architecture and a loss of observable capsular (Bowman’s) space. Hepatic inflammatory responses were largely observed in the portal triad area of tissue sections encompassing the portal vein, hepatic artery, and bile ductule. Also, histopathological evaluation of intestinal infiltrates demonstrated more robust infiltration of the lamina propria and submucosa in LNA control mice. To show that the inflammation was resulting from engrafted human PBMCs and to quantitate the extent of response for statistical comparison, IHC was performed with digital histopathological image analysis to detect and quantify human CD3+ T cells present in the tissue sections. Since our previous work engrafting PBMCs from patients with autoimmune disease into immunodeficient mouse recipients showed that target organ inflammation was comprised of CD4+ and CD8+ T cells [31], IHC for CD3 was used here to analyze both immune cell subtypes collectively. Our results demonstrate statistically significant reductions in CD3+ human T cell infiltrates in the kidney by 17-fold (p = 4.4 × 10^−13^), the small intestine by 2.4-fold (p = 1.2 × 10^−8^), and the liver by 6.2 fold (p = 7.5 × 10^−12^) with LNA-mediated inhibition of miR-21, miR-29a, and miR-29b (**Figure 9**). Spleens of all mice showed high levels of CD3+ human T cell detection, which is indicative of successful human immune cell engraftment and reconstitution *in vivo*. These results establish feasibility of this chimeric model platform to study LNA-based therapeutics to control autoimmune-mediated inflammatory responses and indicate that inhibition of miR-21, miR-29a, and miR-29b can suppress disease pathology.

**Figure 9.**
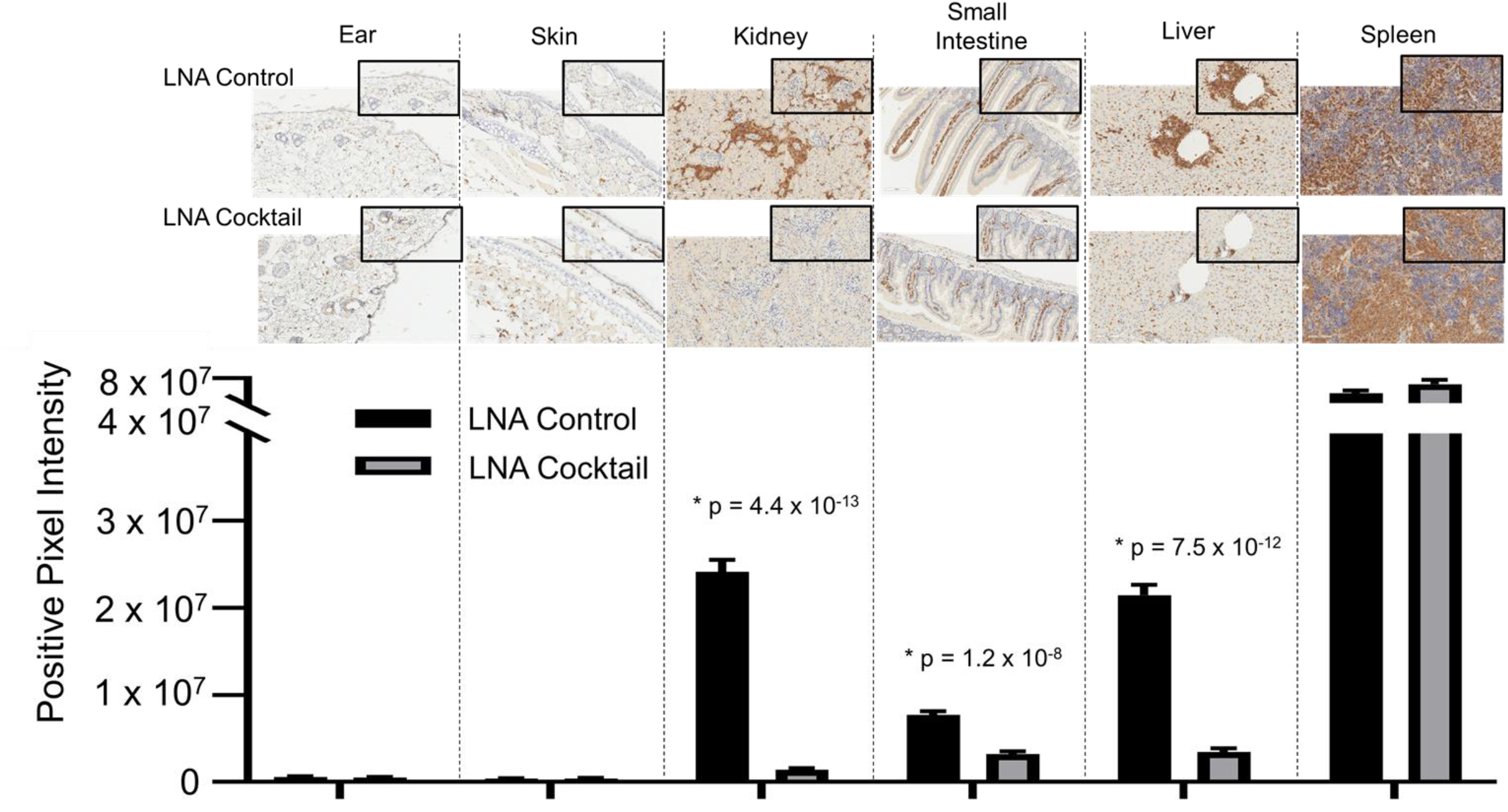
Immunohistochemical analysis of the infiltrates from chimeric mice confirm the presence of human CD3+ T cells in target organs. Target organ inflammation was measured in chimeric mice by IHC for human CD3 detection to compare the effects of a cocktail of locked nucleic acid miR antagonists (LNA cocktail) targeting miR-21, miR-29a, and miR-29b to a nonsense control treatment (LNA control). Large panel images taken at 20X magnification (scale bar representing 100 μm) and insets taken at 40X (scale bar representing 50 μm). Histopathological assessment was performed as described in the materials and methods by digital analysis of scanned slides to evaluate relative CD3 expression. Values are the mean ± SEM with indicated p values shown. *All values of p ≤ 0.05 were considered statistically significant.

## DISCUSSION

Elucidating the pathogenesis and molecular mechanisms driving inflammatory responses in SLE can yield desperately needed diagnostics and targeted therapies. The first approved therapy for lupus was Aspirin (1948), which was followed shortly thereafter by corticosteroids and then hydroxychloroquine in 1955. The next, and first targeted therapy, was not approved until a half century later (2011); belimumab is a recombinant monoclonal antibody targeting B-cell activating factor (BAFF) that is approved for adult patients with active LN. Subsequently, voclosporin (inhibitor of calcineurin-mediated T cell activation approved for adult patients with LN) and Anifrolumab-fnia (monoclonal antibody inhibitor of type I interferon receptor approved for adult patients with moderate to severe SLE) were both approved in 2021. Consequently, despite 75 years of research, only 2 targeted agents (belimumab and anifrolumab-fnia) and several non-specific systemic anti-inflammatory agents have been approved as prophylaxis for this autoimmune disease. Similarly, clinical tests to diagnose SLE or to predict flares have also lagged behind other scientific advances in medicine. In the “omics” era of biomarker discovery via DNA/RNA sequencing, epigenetics, and proteomics, SLE is still largely diagnosed and treated according to clinical classification criteria, as defined by the American College of Rheumatology (ACR) and the Systemic Lupus International Collaborating Clinics (SLICC) [39, 40]. Typical laboratory testing conventionally ordered in the clinic to aid in SLE diagnosis includes anti-nuclear antibodies (ANA), anti-double-stranded DNA antibodies (anti-dsDNA), and complement levels, but these tests have limited sensitivity and specificity [41-43]. While the gold standard for LN diagnosis is a kidney biopsy to confirm renal involvement, this procedure is invasive and has associated risks and complications. In this study, we identify a novel exosome-mediated miR signaling pathway that is associated with SLE disease activity and demonstrate that blocking upregulated miRs in exosomes can inhibit the pathogenic inflammatory response. SLE-associated miRs packaged in exosomes can potentially be developed as clinical/companion diagnostics and miR antagonists can be a novel area of future therapeutic development.

Collectively, the results from this study suggest a putative mechanism of estrogen-mediated miR synthesis contributing to exosome-encapsulated miR signaling and induction of inflammatory responses in SLE by signaling through TLR7 or TLR8 (**Figure 10**). In this pathway, estrogen stimulates the production of miRs by inducing the expression of RNA polymerase III, Dicer1, AGO2, and Drosha. Intracellular miRs are packaged and secreted in exosomes for delivery to recipient cells. The miRs within exosomes act as cytokines to mediate cell-to-cell communication in the inflammatory response; following this nomenclature, we have previously referred to exosome-encapsulated miRs as miRokines [36]. Upon entry into the new cell, miRokines can function in the canonical pathway by associating with RNA-induced silencing complexes (RISC) to bind complementary mRNAs and suppress gene expression or non-canonically by binding to TRL7 or TLR8. Activation of TLR7 and/or TLR8 by miR ligands can stimulate additional exosome secretion and proinflammatory gene expression, which can contribute to SLE pathogenesis.

**Figure 10.**
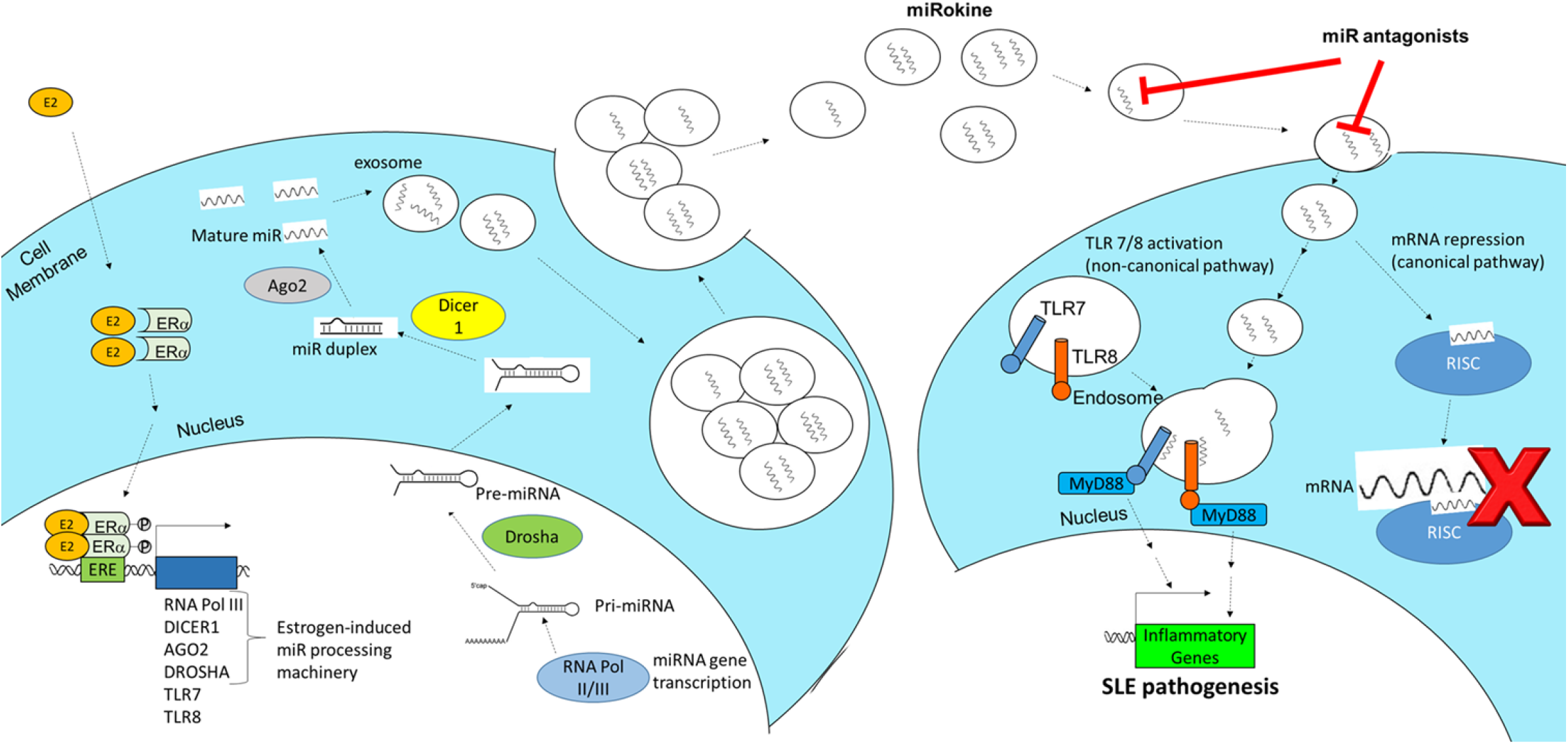
Schematic of the proposed mechanism of estrogen-mediated miR production and inflammation via extracellular vesicle signaling in SLE. Estrogen (E2) enters an immune cell, dimerizes with estrogen receptor (ER)α, translocates to the nucleus, and promotes the expression of miRNA (miR) processing machinery, including RNA polymerase III, Dicer1, AGO2, and Drosha. These miRs are packaged and secreted in EVs, which are taken up by recipient immune cells. When EV-derived miR cargo is taken up, it can function to regulate gene expression in the recipient cell via two pathways. The canonical pathway involves miRs binding to target mRNAs through the RNA-induced silencing complex (RISC). In the non-canonical pathway, EV-encapsulated miRs fuse with endosomes and bind to TRL7 or TLR8 to stimulate proinflammatory gene expression and additional EV secretion, which promotes SLE pathogenesis.

Estrogen has a pleotropic effect on immune system function and can significantly stimulate macrophage and lymphocyte proliferation [44]. Our previous studies have shown that estrogen can lower the threshold of immune cell activation and that this response is more robust in females [21]. These data suggested that estrogen may enhance immune system activation, which could augment infectious disease responses, but also may contribute to autoimmune disease development and pathogenesis. In SLE, disease-associated sex-bias is evident [45], as most clearly demonstrated by the patient population; the adult premenopausal female to male ratio of SLE is 9:1, which then becomes closer to 2:1 either during childhood or post menopause [46]. Here, we discovered that estrogen can stimulate the expression of proteins involved in intracellular miR processing. While a direct correlation of estrogen and exosome production has not been established, previous studies have demonstrated that estrogen can have a significant impact on the intracellular regulation of miR expression. Estrogen receptor (ER)α positive and hormone responsive breast cancer cell lines (MCF-7 and ZR-75.1) differentially regulated 172 miRs with estrogen treatment [47]. Moreover, unique miR expression profiles have been reported in human breast cancer and these differences correlated with expression of ER, tumor stage, number of positive lymph nodes, and vascular invasion (reviewed in [48]). Our results demonstrate that estrogen stimulation of primary human PBMCs can upregulate miR processing machinery, which may lead to the unique intracellular miR profiles observed in other studies and account for the distinct miRs packaged within exosomes of SLE patients.

Our data characterize EVs and the associated RNA biocargo in addition to evaluating the therapeutic strategy of miR inhibition in SLE. While the delivery of RNA biocargo within EVs to influence recipient cell function has only recently been explored therapeutically, the clinical success of SARS-CoV-2 mRNA vaccines has validated this as a biologically relevant and viable therapeutic strategy. Both the mRNA-1273 and BNT162b2 mRNA-based vaccines for SARS-CoV-2 are packaged into lipid nanoparticles (LNPs) that are approximately 100 nm in size [49]. The LNPs are taken into recipient cells by endocytosis and the mRNA biocargo is translated into SARS-CoV-2 Spike protein, which ultimately elicits a protective immune response targeting this viral antigen. Similarly, both the liposomes used here to deliver LNA antagonists and the exosomes used for RNA sequencing analysis from plasma samples were also approximately 100 nm in size. While blocking RNA expression is a relatively new therapeutic strategy, there have been four small interfering (si)RNA drugs approved by the FDA to date: patisiran, givosiran, lumasiran, and inclisiran [50]. Interestingly, patisiran is a double stranded siRNA that is delivered via encapsulation within LNP delivery vectors. LNAs represent the third generation of chemically modified RNA therapeutics and exhibit binding affinities and RNA neutralization capacities far superior to siRNA [51]. In addition to enhanced affinity, potency, and specificity to RNA targets, LNAs also exhibit bioavailability and stability, as indicated by the diverse tissue distribution and long half-life *in vivo* observed in the clinical trials of a miR-122 LNA antagonist (miraversen) [52]. Considering the preclinical and clinical development of RNA targeting and delivery through LNPs similar in size to exosomes, our data suggests that exosome-encapsulated RNAs are biologically significant and represent a novel pharmacological platform to evaluate therapeutically in SLE.

Despite the biological relevance of exosomal biocargo signaling in regulating inflammatory processes and the potential of exosomes to be used as disease-associated biomarkers, very little data has been generated to date in the study of autoimmune disease. While distinct miR signatures have been identified in exosomes derived from LN patients and specific miRs have been associated with renal fibrosis and disease activity [19, 53, 54], very few studies have evaluated blood derived exosomes in SLE. Moreover, although these studies of exosomes in SLE blood circulation identified differential regulation of specific miRs (miR-145a, miR-21, and miR-155), the EVs evaluated and data generated may be physiologically irrelevant considering that serum sources are subject to platelet-derived exosome secretion/contamination while the blood sample is clotting during experimental processing [55]. Our results demonstrate plasma-derived EVs are upregulated in both inactive and active SLE patients and contain distinct small RNA profiles. Interestingly, vault (v)RNAs (*VTRNA1-1 and VTRNA1-2*) were significantly downregulated in active LN patients. vRNAs have been detected in exosomes previously and are thought to play a role in intracellular and nucleocytoplasmic transport processes [56].

Furthermore, previous research has shown that vRNAs can confer resistance to autophagy when overexpressed [57, 58] as well as inactivate protein kinase R signaling, which inhibits interferon responses [59]. Consequently, these findings suggest that exosome delivered vRNAs could have significant influence over the cellular regulation of autophagy [60] and interferon signaling [61] that is observed in patients with active LN.

In summary, the results of this study propose a novel estrogen-mediated pathway of intracellular miR production and demonstrate dysregulated RNA expression patterns in EVs of SLE patients that potentially can be targeted therapeutically to suppress disease pathogenesis. Future work will be directed toward defining the precise mechanistic role of estrogen in this process, examining the candidacy of exosomes as disease-associated diagnostic biomarkers, and identifying additional miRs to target therapeutically. In addition to miRs, exosome biocargo includes surface-expressed and intra-exosomal protein, mRNA, vRNA, and long noncoding (ln)RNA. The roles of these additional RNA species in cell-to-cell communication and as putative biomarkers have yet to be explored in SLE. Furthermore, biomarker characterization of exosome protein expression can define cell of origin, which may be used to predict specific organ involvement early in the disease course. Previous studies have also demonstrated that exosomes can contain cytokines [62]; however, the biological relevance of this observation and the potential role that this may play in immunopathology has not been described. Through these follow-up studies, the roles of different exosomal biocargo in the diagnosis and treatment of SLE will be better characterized.

## ACKNOWLEGEMENTS

This research was supported by the Clinical Research Center/Center for Clinical Research Management at Ohio State University’s Wexner Medical Center and The Ohio State University College of Medicine under protocol IMMUN39: Ohio State University Rheumatology and Immunology Biorepository (CCTS ID#: 4735). We would also like to extend our gratitude to the volunteers that participated in this study and would like to acknowledge Jeff Hampton and Giancarlo Valiente for their contributions to the data collection and analysis portion of this work while working at The Ohio State University Wexner Medical Center.

## COMPETITING INTERESTS

None declared.

## DATA SHARING STATEMENT

All of the data collected during this study, the data analysis plan, and the treatment protocol are on-file for sharing immediately following publication to anyone who wishes to access the data for any purpose with no end date. Proposals should be directed to. To gain access, requestors will need to sign a data access agreement.

## PROVENANCE AND PEER REVIEW

Not commissioned; externally peer reviewed.

## ETHICAL APPROVAL AND CONSENT FOR PUBLICATION

Human studies were approved through The Ohio State University Wexner Medical Center (OSUWMC) Institutional Review board (IRB) Protocol Numbers: 2011H0094, The Ohio State University Rheumatology and Immunology Lupus Clinic Registry; and 2015H0340, Multi-Source Examinations of Biomarkers in Autoimmune Diseases. Animal study design and protocol 2017A00000032 was approved by the Institutional Animal Care and Use Committee and University Laboratory Animal Resources at OSUWMC.

## AUTHOR CONTRIBUTIONS

Conceived and designed the experiments of the study: (NY, ER, WJ). Data collection and analysis: (NY, ES, KJ, ER, WJ). Performed experiments: (NY, ES, KJ, ER). Edited manuscript: (NY, ES, KJ, LW, ER, WJ). Statistical assessments: (NY, ER). Wrote manuscript: (NY, ER, WJ). Contributed reagents/materials/analysis tools: (LW, ER, WJ). Made substantial, direct and intellectual contribution to the work, and approved it for publication: (NY, ES, KJ, LW, ER, WJ).

